# Evolutionary rescue accelerates competitive exclusion in a parasite community

**DOI:** 10.1101/2025.09.25.678511

**Authors:** Samuel TE Greenrod, Daniel Cazares, Weronika Ślesak, Tobias E Hector, R. Craig MacLean, Kayla C King

## Abstract

Environmental stress drives biodiversity loss by altering competitive hierarchies and pushing taxa towards extinction. Parasites and their communities are particularly vulnerable to stress due to environmental sensitivity of infection steps, variation in species tolerance during co-infections, and dependence on host fitness. Parasite populations might avoid extinction through evolutionary rescue – whereby rapid adaptation to stress enables persistence – but whether this process can preserve community diversity remains unclear. Here, we study the impact of evolutionary rescue in a simple parasite community by propagating populations of two viral parasites (bacteriophages ϕ14-1 and ϕLUZ19) of *Pseudomonas aeruginosa* in monoculture and co-culture under two thermal conditions, a control temperature (37°C) and a high temperature that restricts ϕ14-1 growth (42°C). We show that evolutionary rescue of ϕ14-1 prevented extinction in monoculture. Rescue of this phage in co-culture made it a superior competitor, and it replaced ϕLUZ19 as the dominant phage at high temperature. We determine that evolutionary rescue occurred through mutations in genes linked to attachment to bacterial hosts and within-host replication. We also show that competitive suppression by ϕ14-1 constrained ϕLUZ19 molecular evolution. Our findings suggest that evolutionary rescue can prevent the extinction of some parasites, but may inadvertently destabilise the community and facilitate further biodiversity loss. This work underscores the need to take an eco-evolutionary approach to predict the responses of communities to global climate change.

## Introduction

Environmental stress is a primary driver of biodiversity loss (1,2). On an ecological timescale, only those species with environmental optima and tolerance ranges best aligned with prevailing conditions can persist (3–5). As community composition shifts towards fewer species, the probability of extinction events (6) and ecological tipping points (7) is heightened. On an evolutionary timescale, however, biodiversity could be maintained via evolutionary rescue, whereby adaptation to environmental stress facilitates population recovery (7,8). Evolutionary rescue has been shown to prevent diversity loss in communities and increase the prevalence of rare taxa whose relative fitness increases following adaptation (9–11). Alternatively, evolutionary rescue can alter the competitive hierarchy in communities and drive competitor decline and exclusion (12,13). Whether evolutionary rescue can counteract the diversity-eroding impacts of environmental stress is unclear. Understanding how environmental stress affects biodiversity requires consideration of both ecological and evolutionary dynamics in communities (14).

Parasite communities are expected to be among those most disrupted by climate change, with consequences for global disease dynamics and ecosystem stability (15). Parasite diversity has been shown to decrease with thermal stress through widening inter-parasite fitness differences (16) and increasing host resistance (17). Diversity can also be lost through thermal alterations to parasite life-history strategies (18) or a reduction in niche differences (19,20). Microbial parasites offer a powerful model system to study evolutionary rescue in communities due to their rapid evolution rates and frequent competitive interactions with other parasites (21,22). Parasites can be highly sensitive to thermal extremes ((23–25), Greenrod et al., in press) and, in isolation, have been shown to avoid thermal extinction through evolutionary rescue (16,26). However, the evolutionary responses to thermal stress in communities are more complex. Inter-parasite competition can constrain evolutionary rescue by reducing population growth rates and mutational supply (5,18,27,28). Alternatively, rescue may be promoted with competition and thermal stress selecting on the same parasite life-history traits ((29), Greenrod et al., in press). Evolutionary rescue may thus alter parasite competitiveness and increase the risk of competitor exclusion (13,30,31) in response to stressful temperatures.

Evolutionary rescue is predicted to increase the absolute fitness of a parasite species during warming and prevent extinction. While competition in parasite communities could constrain evolution through limiting mutational supply (28), adaptation could skew the competitive hierarchy towards the rescued, thereby accelerating competitive exclusion. To test these predictions, we passaged two lytic viral parasites (thermal generalist ϕLUZ19 and specialist ϕ14-1) through evolutionarily static populations of a bacterial host, *Pseudomonas aeruginosa* (Fig. 1). These phages are obligate killers; following phage attachment and infection of host cells, they immediately initiate replication and host cell lysis (death). Phages were evolved under a control temperature (37°C) or high temperature that restricts the growth of the ϕ14-1 phage (42°C) (32) in monoculture or co-culture. We tested for effects on phage growth rates and competitiveness by conducting phenotypic assays of phage infectivity in the absence and presence of a phage competitor. We also used sequencing to measure changes in the genomic composition of phage populations through time.

**Figure 1.**
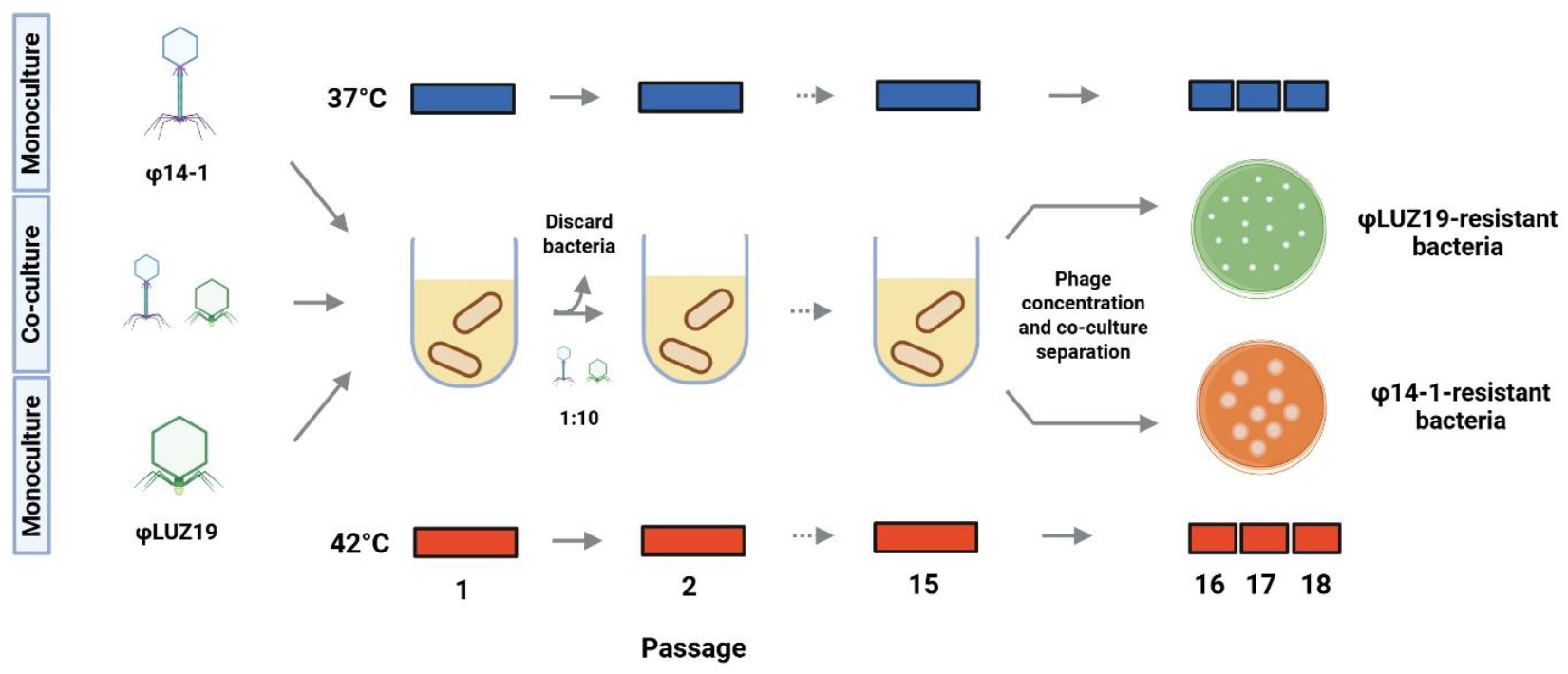
Overview of phage evolution experimental framework. Phages evolved in monoculture or co-culture at 37°C or 42°C for 15 passages. At the end of each passage, phages were isolated from lysates and used to infect fresh, evolutionarily static bacterial hosts (Passages 1-15). At the end of the selection experiment, phages were concentrated and co-cultures separated through three rounds of confluent lysis plating on phage-resistant bacteria (Passages 16-18) under the same thermal regime as earlier passages. Phage icons illustrate the two different phages used in the experiments (ϕ14-1, myovirus in blue; ϕLUZ19, autographivirus in green) (33) and are used hereafter to refer to phages in figures. Figure was created using BioRender.

## Results

### Evolutionary rescue prevents phage extinctions in monoculture

While the ancestral ϕLUZ19 grows equally at 37°C and 42°C, ancestral ϕ14-1 populations show no signs of growth at 42°C at a starting density of 10^4^ PFU/ml (Fig. 2A). Passaging at this density would have thus resulted in dilution to extinction. We hypothesised that the thermally sensitive phage ϕ14-1 would avoid extinction by rapidly adapting to tolerate thermal stress. While ϕ14-1 growth was initially restricted at 42°C, it rapidly reached and maintained high densities (>10^9^ PFU/ml) across temperatures (Fig. 2B). High densities were also reached by the thermal generalist ϕLUZ19. Across both phages, no replicate populations went extinct.

**Figure 2.**
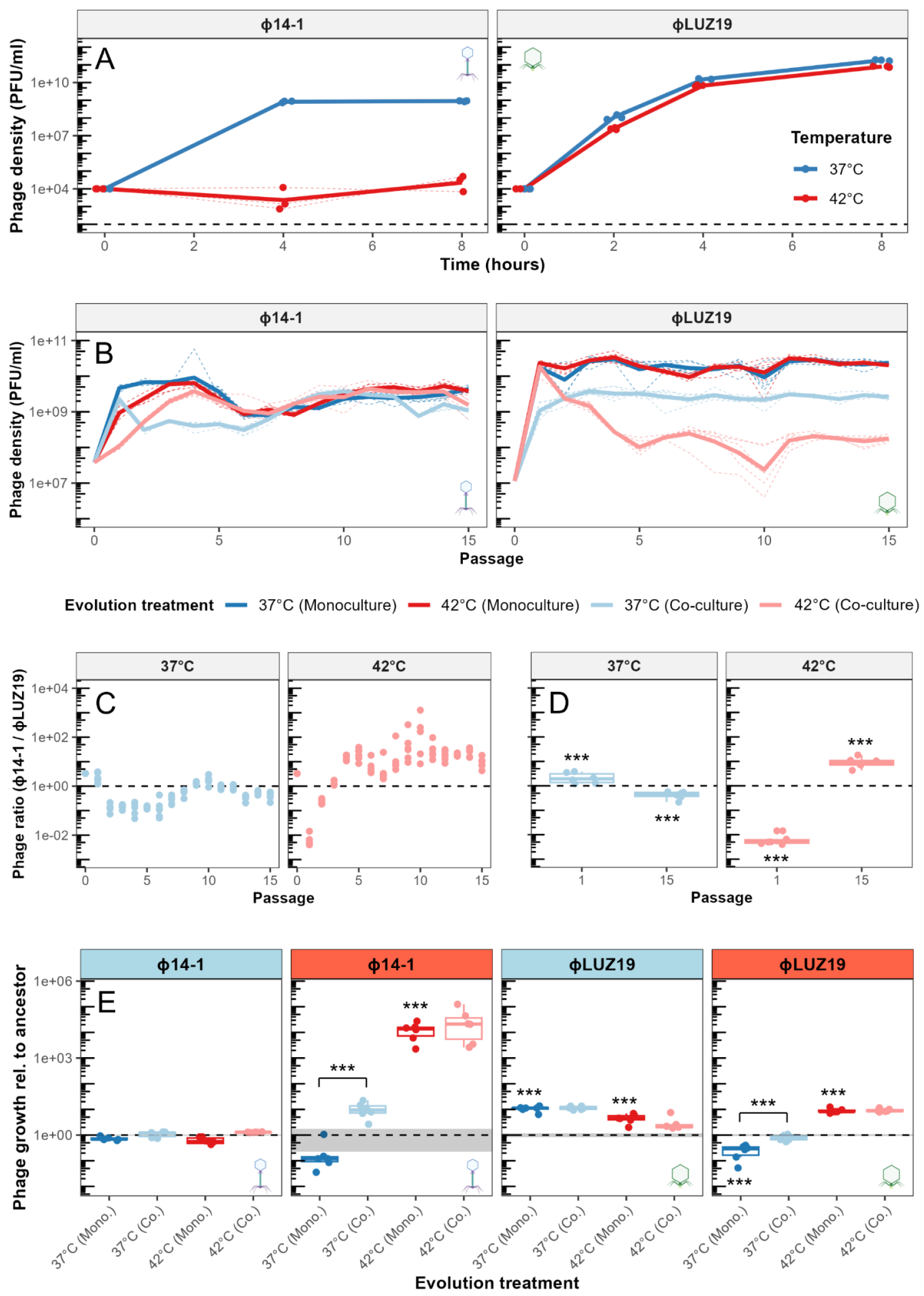
Evolutionary rescue alters the competitive hierarchy. **A**) Growth curves of ancestral phages across temperatures. Lines show ancestral phage growth curves at 37°C (deep blue) and 42°C (deep red). Black dashed line marks the limit of detection. **B**) Phage population densities under each evolution treatment across passages. Lines show phage population densities during the first 15 evolutionary passages of phages in monoculture (37°C in deep blue, 42°C in deep red) and in co-culture (37°C in light blue, 42°C in light red). Solid lines show average values of six biological replicates (each shown as dashed line). Values show densities at the end of each passage prior to dilution. **C**) Ratio of ϕ14-1 to ϕLUZ19 densities in co-culture treatments across passages. Values show phage ratios at the end of each passage where each dot shows a replicate population. Phage ratio of 1:1 is shown with black dashed line. **D**) ϕ14-1 to ϕLUZ19 ratios in co-culture treatments at passage 1 and passage 15. Asterisks show significant differences to a phage ratio of 1:1 (black dashed line) where *** = p < 0.001. **E**) Growth rates of end-point evolved phage populations relative to the ancestral population tested in monoculture at 37°C (light blue strip) and 42°C (light red strip). ϕ14-1 populations were compared after 4h growth and ϕLUZ19 populations were compared after 2h growth. Evolved phage growth (box) is shown relative to ancestor (black dotted line). Shaded grey region shows ancestor standard errors from three biological replicates, each being an average of three technical replicates. Asterisks above line connectors show significant differences between evolved populations. Asterisks above boxes show significant differences from the ancestor. *** = p < 0.001. Co-culture lines were not compared to the ancestor. Otherwise, no asterisk reflects non-significance.

We measured bacterial growth using optical densities at 37°C and 42°C (Fig. S1) to account for any host-mediated variation in phage densities. Without phage, *P. aeruginosa* had significantly higher growth rates at 42°C compared to 37°C (F_3,62_ = 78.9, p < 0.001). Based on a standard curve of optical density to colony forming units (Fig. S2), average bacterial densities at 42°C were found to be approximately double those at 37°C across an 8h passage. Yet, phage densities at each passage were the same or lower at 42°C than 37°C indicating that variation in phage densities could not be explained by differences in bacterial growth rates.

To determine whether ϕ14-1 densities were maintained at 42°C through evolutionary rescue, we conducted population growth assays for the end-point evolved phage populations at 37°C and 42°C (Fig. 2E). Phage population growth rates depended on an interaction between evolution treatment (37°C or 42°C) and assay temperature (ϕ14-1: F_2, 19.1_ = 199, p < 0.0001; ϕLUZ19: F_2,19.1_ = 100.0, p < 0.0001). ϕ14-1 lines evolved at 42°C showed a significant increase in growth at 42°C compared to ancestor (t(20.9) = −18.1, p < 0.0001). However, 37°C evolved populations showed no change in growth at 37°C (t(20.9) = 0.64, p = 0.987), likely due to phages reaching close to carrying capacity at the point of measurement (Fig. S3). ϕLUZ19 lines also had significantly higher growth rates at their evolved temperatures compared to the ancestor (37°C: t(20.9) = −7.68, p < 0.0001; 42°C: t(20.9) = −7.04, p < 0.0001).

Higher growth rates under evolved conditions may reflect merely adaptation to the host rather than temperature-specific fitness changes. We assessed phage growth at allopatric (e.g., mismatched) temperatures (Fig. 2E). ϕ14-1 evolved populations showed no improvement in growth at allopatric temperatures (37°C evolved populations: t(20.9) = 2.37, p = 0.212, 42°C evolved populations: t(20.9) = 1.00, p = 0.912). In addition, ϕLUZ19 lines evolved at 37°C exhibited a significant growth rate decrease at 42°C (t(20.9) = 5.13, p < 0.001). However, ϕLUZ19 lines evolved at 42°C were found to also have significantly higher growth rates at 37°C than ancestor (t(20.9) = −4.79, p < 0.01) indicative of adaptation to the host as opposed to the thermal regime in this instance.

### Evolutionary rescue alters the competitive hierarchy

While thermal adaptation occurred in monoculture, we hypothesised that adaptation would be restricted in co-cultures due to evolutionary constraint from inter-phage competition (5,27). Both ϕ14-1 and ϕLUZ19 were initially found to have lower population densities in co-culture than in monoculture (Fig. 2B). At 37°C, ϕ14-1 and ϕLUZ19 had ∼10-fold lower densities in co-culture up to passage 6, after which ϕLUZ19 densities remained suppressed and ϕ14-1 densities converged with those of the monoculture populations. While ϕ14-1 was heavily restricted by ϕLUZ19 in early passages at 42°C, it reached similar densities to monoculture lines by passage 4. In contrast, ϕLUZ19 co-culture densities were initially high at 42°C but rapidly decreased before stabilising.

We hypothesised that the population decline in ϕLUZ19 42°C co-culture populations occurred due to a shift in the competitive equilibrium with ϕ14-1 following evolutionary rescue. We assessed the competition dynamics by tracking the ratio of ϕ14-1 and ϕLUZ19 densities across co-culture passages where a ratio > 1 reflects a ϕ14-1 competitive advantage and vice versa. At 42°C, the ratio of ϕ14-1 densities relative to ϕLUZ19 fell in the initial passage but then rapidly increased before stabilising at passage 5 (Fig. 2C). The phage ratio conversely remained relatively stable across all passages in the control. By analysing phage ratios at passages 1 and 15, we found that, at 42°C, ϕ14-1 had a significant competitive disadvantage at passage 1 (t(19.9) = −27.5, p < 0.001) but a significant advantage by passage 15 (t(19.9) = 11.7, p < 0.001). ϕ14-1 competitive advantage showed a small decrease between passage 1 and 15 in the control (Passage 1: t(19.9) = 3.9, p < 0.001; Passage 15: t(19.9) = −4.7, p < 0.001). We further investigated the competitive profile of the rescued phage in time-shifted, direct competition assays at low and high MOI between evolved ϕ14-1 co-culture populations against a ϕLUZ19 ancestral population (Fig. S4). At low MOI, the restriction of ϕLUZ19 ancestral growth by ϕ14-1 was significantly lower in the ϕ14-1 37°C co-culture population than the ϕ14-1 ancestor at 42°C (t(21.6) = −4.4, p < 0.01). At high MOI, ϕLUZ19 ancestor restriction at 42°C was significantly greater in the ϕ14-1 42°C co-culture populations than the ϕ14-1 ancestor (t(21.6) = 3.9, p < 0.01), but the magnitude of change was relatively small. The ϕ14-1 competitive advantage in 42°C co-culture populations did not reflect an escalating increase in competitive fitness across evolutionary time.

By restricting phage growth, we hypothesised that the presence of a competitor would constrain phage thermal adaptation. We assessed thermal adaptation by comparing phage thermal phenotypes in monoculture and co-culture evolved populations (Fig. 2E). For both phages, we found that growth rates depended on an interaction between evolution treatment (monoculture and co-culture) and assay temperature (ϕ14-1: F_4,39_ = 132, p < 0.0001; ϕLUZ19: F_4,39_ = 108, p < 0.0001). There was no significant difference in growth rates between monoculture and co-culture evolved populations at their evolved temperatures (37°C − ϕ14-1: t(39) = −0.87, p = 0.99; ϕLUZ19: t(39) = 0.38, p = 1.0; 42°C − ϕ14-1: t(39) = −1.1, p = 0.98; ϕLUZ19: t(39) = −0.19, p = 1.0). Surprisingly, phages evolved at 37°C in co-culture showed significantly higher fitness at 42°C than those evolved in monoculture (ϕ14-1: t(39) = −9.19, p < 0.0001; ϕLUZ19: t(39) = −6.04, p < 0.0001).

### Temperature and competition select for mutations in tail proteins and replication machinery

To further understand the genomic causes of adaptation to temperature and competition, we conducted phage population sequencing and determined the identity and frequency of genetic variants (Fig. 3; Table S1). Putative adaptive variants were defined as those with a frequency > 20% and which occurred in genes which acquired mutations in at least two biological replicates across all treatments (Table S2).

**Figure 3.**
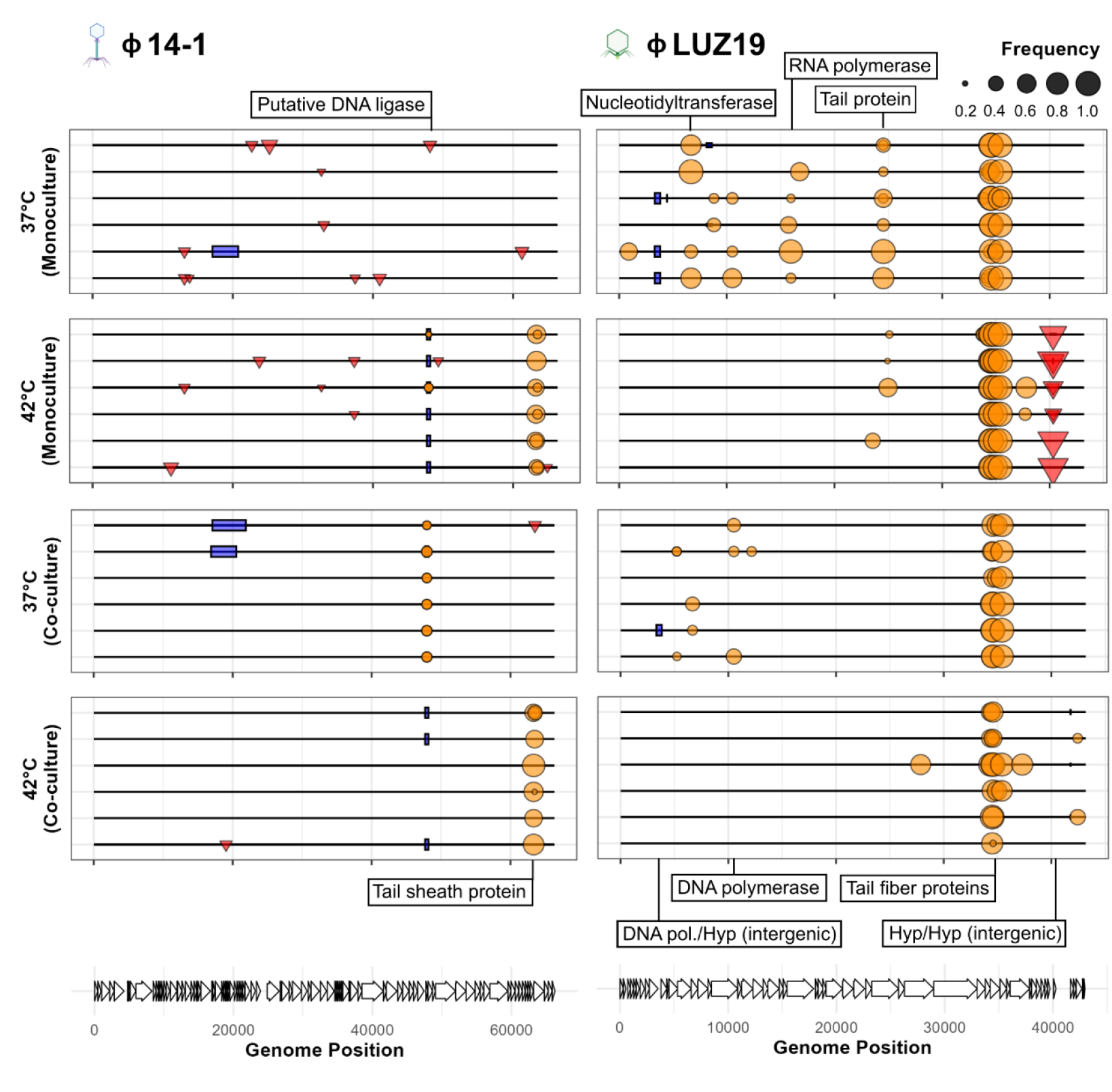
Competition and temperature select for mutations in tail proteins and replication machinery. Mutation plots show genetic variants associated with thermal adaptation and competition in phage lines. Lines represent individual biological replicates. Symbols within plots show variants across the phage genome at >20% prevalence and which were not observed in the ancestral population. Symbols reflect mutation type where circle = SNP, box = deletion, inverted triangle = insertion. Length of deletion bars represent the size of deletion except for the ϕ14-1 deletion at ∼48kb which is a 1bp deletion but given a fixed size for visibility. Labels show annotations for genes which contain mutations in at least three replicate populations in the same evolution treatment, reflecting parallel evolution. All putative adaptive variants are presented in Table S2. Putative DNA ligase in ϕ14-1 was originally annotated a hypothetical protein but has high homology to Pseudomonas phage PhL_UNISO_PA-DSM_ph0031 DNA ligase protein.

As ϕLUZ19 growth is restricted by slow host attachment rates (32), we hypothesised that ϕLUZ19 mutations would occur in tail proteins responsible for attachment. The most prominent genetic changes in the ϕLUZ19 evolved populations were a series of high frequency SNPs in two genes encoding a tail protein and tail fiber protein. While the tail protein mutations were only found in monoculture lines, tail fiber protein mutations were found across all temperature and phage combination treatments suggesting that these reflect adaptation to the host rather than to inter-specific competition or thermal stress. ϕLUZ19 42°C evolved populations also all contained insertions in an intergenic region between two hypothetical proteins, suggesting that altered regulation of gene expression may also contribute to thermal adaptation in ϕLUZ19. Other mutations were exclusively found in 37°C evolved populations and included putative bacterial immune system-associated nucleotidyltransferase (34), RNA polymerase, and DNA polymerase mutations. These mutations may contribute to ϕLUZ19 low temperature adaptation by increasing phage replication within cells.

We also sequenced the ϕ14-1 evolved populations to confirm that the changes to thermal phenotypes reflect evolutionary rescue rather than a plastic response. ϕ14-1 rescue at 42°C was linked to a series of high frequency SNPs in a gene encoding the phage tail sheath, a phage component involved in DNA transfer into bacterial cells (35). We also observed parallel deletions and SNPs in the 42°C monoculture and 37°C co-culture populations in a putative DNA ligase (BlastP: 97.47% identity, 95% sequence overlap with Pseudomonas phage PhL_UNISO_PA-DSM_ph0031 DNA ligase protein), a protein that is essential for phage DNA replication and fitness (36,37). These results imply that competition and high temperature co-select for altered DNA replication in ϕ14-1.

### Thermal stress and competition shape phage molecular evolution

Phage populations evolved at high temperatures differed from control lines both in terms of thermal phenotypes and the mutations they acquired. We hypothesised that 37°C and 42°C evolved populations would also diverge at the whole-genome level due to differing evolutionary trajectories. PCoA analysis based on phage Euclidean genetic distances showed that both ϕ14-1 and ϕLUZ19 had significant genetic divergence between 37°C and 42°C evolved populations (ϕ14-1: ANOSIM: R = 1.0, p < 0.01; ϕLUZ19: ANOSIM: R = 0.91, p < 0.01) (Fig. 4A). We then hypothesised that evolution rates would be highest in the lines that experienced the greatest change in growth rates. ϕ14-1 had significantly higher evolution rates at 42°C than at 37°C (F_1,10_ = 30.3, p < 0.001) (Fig. 4B). No significant difference in evolution rates was observed between temperatures for ϕLUZ19 populations (F_1,10_ = 1.2, p < 0.30).

**Figure 4.**
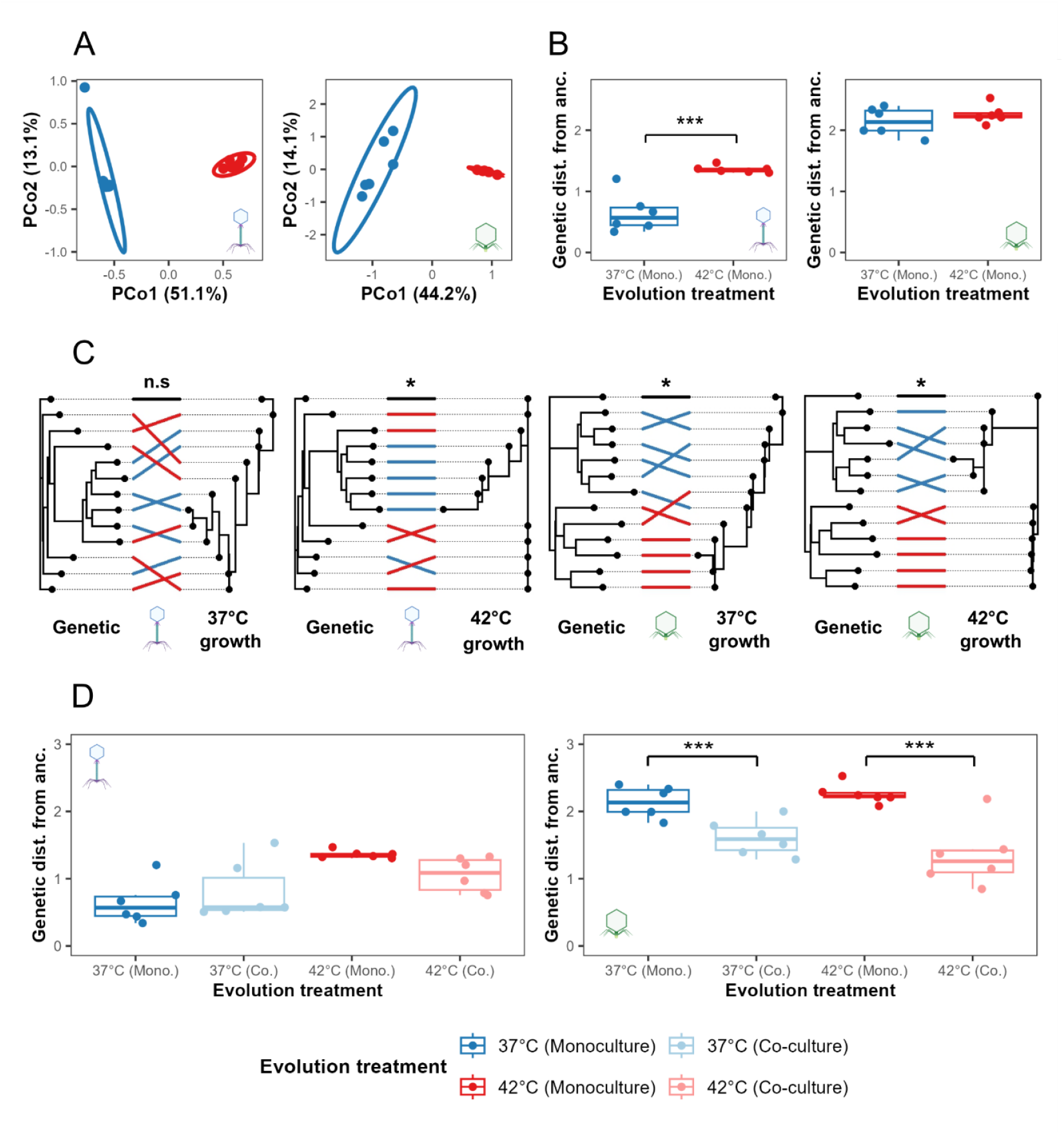
Thermal stress accelerates and competition constrains phage molecular evolution. **A**) Genomic divergence between evolved monoculture phage populations. PCoA plots show Euclidean genetic distance clustering between 37°C and 42°C evolved phage populations based on mutation position and frequency. **B**) Evolution rates of evolved monoculture phage populations based on the Euclidean genetic distance from ancestor. *** = p < 0.001. No asterisk reflects non-significance. **C**) Congruence analysis of neighbour-joining trees constructed based on Euclidean genetic distances (left-hand plots, labelled “Genetic”) and Euclidean distances based on phage growth rates at 37°C and 42°C (right-hand plots, labelled “37°C growth” and “42°C growth”, respectively). Congruence is shown by the alignment of tips corresponding to individual replicate populations between trees. High congruence is shown by few cross-overs. 37°C monoculture population connections are shown in deep blue, 42°C monoculture populations in deep red, and the ancestral phage in black. Trees are rooted using the ancestral phage. **D**) Evolution rates of evolved monoculture phage populations compared to evolved co-culture populations. Evolution rates are determined based on Euclidean genetic distance from ancestor. *** = p < 0.001. No asterisk reflects non-significance.

As phages showed both phenotypic and genomic divergence based on thermal regime, we hypothesised that genetically similar phage populations may have similar thermal fitness. We assessed the relationship between thermal fitness and genomic change by measuring congruence between neighbour-joining trees constructed based on Euclidean phenotypic and genetic distances (Fig. 4C). Highly significant congruence was observed for both phages based on growth rates at 42°C (ϕ14-1:M^2^_xy_ = 5.8, p < 0.01; ϕLUZ19:M^2^_xy_ = 15.9, p < 0.001). For 37°C growth, significant congruence was observed for ϕLUZ19 (M^2^_xy_ = 20.4, p < 0.05) but not for ϕ14-1 (M^2^_xy_ = 8.1, p = 0.18).

Given the evolutionary constraint of competition, we also hypothesised that co-culture evolved populations would diverge from monoculture evolved populations and have slower evolution rates. Significant genomic divergence was observed between monoculture and co-culture populations for ϕ14-1 at 37°C and for ϕLUZ19 at 42°C (ϕ14-1: ANOSIM: R = 0.18, p < 0.005; ϕLUZ19: ANOSIM: R = 0.81, p < 0.005). Divergence was not significant for ϕ14-1 at 42°C or for ϕLUZ19 at 37°C (ϕ14-1: ANOSIM: R = 0.27, p = 0.057; ϕLUZ19: ANOSIM: R = 0.15, p = 0.07) (Fig. S5). ϕ14-1 populations had similar evolution rates in monoculture and co-culture (37°C: t(20) = −0.972, p = 0.77; 42°C: t(20) = 1.77, p = 0.31). However, ϕLUZ19 evolved populations had significantly slower evolution rates in co-culture compared to monoculture at both 37°C and 42°C (37°C: t(20) = 3.1, p < 0.05; 42°C: t(20) = 5.3, p < 0.001) (Fig. 4D).

## Discussion

Environmental stress reduces biodiversity by restricting the growth of sensitive community members and destabilising competitive hierarchies (28). We show that the ϕ14-1 phage can avoid heat-driven extinction through evolutionary rescue, even in the presence of a thermally tolerant ϕLUZ19 phage competitor. We further found that competition at permissive temperatures, where both phages grow efficiently, can drive the evolution of elevated ϕ14-1 thermal tolerance. These findings support previous studies by demonstrating that phages are highly evolvable in response to environmental stress (16,38,39). The results nevertheless contradict findings that competitive interactions constrain environmental adaptation by reducing growth rates and mutational supply (5,27). One potential explanation is that, while strong competition may constrain evolution, weak or moderate competition may have increased the strength of selection for ϕ14-1 thermal adaptation (4,28,40). Alternatively, adaptation may have been driven through co-selection by competition and temperature for the same traits. We identified mutations in the same genes in ϕ14-1 populations evolving under both high temperature selection and at permissive temperatures in the presence of competition. The evolution of stress tolerance in free-living organisms has historically been associated with trade-offs in competitive fitness (41–43). Increased selection or mutualistic pleiotropy could mean that competition in parasite systems leads to trade-ups rather than trade-offs with adaptation to environmental stress.

Evolutionary rescue has largely been thought to maintain biodiversity by preventing taxa from becoming extinct (8). Within communities, however, we found that evolutionary rescue can cause one species to become a superior competitor, thereby promoting the competitive exclusion of others. Evolutionary rescue may thus be insufficient to prevent, and may even facilitate, biodiversity loss under environmental stress. Evolutionary rescue can come at an ecological cost where increased environmental tolerance leads a trade-off with growth rates (44,45). However, ecological de-stabilisation can occur via trade-ups if growth rates increase in the recently adapted population (12,13). Competitive shifts may essentially depend on how rescue affects species absolute fitness. We showed that while the rescued phage at 42°C gained a competitive advantage over its sympatric competitor, rescued phage competitiveness did not escalate across evolutionary time and was not elevated against the ancestral competitor. These findings could be explained by the competitor experiencing a decrease in relative fitness following 42°C adaptation (44,45). Competitive dominance may alternatively be specific to co-evolving competitors. By reducing competitor population densities, evolutionary rescue had the additional impact of constraining competitor evolution rates and restricting the acquisition of putative adaptive mutations. Rescue may therefore cause community instability by both depressing competitor population densities and limiting community adaptability in response to future environmental stress (5).

Despite being suppressed by the newly dominant ϕ14-1, ϕLUZ19 competitor phage populations ultimately stabilised at reduced densities. Modern co-existence theory states that co-existence can occur through stabilising mechanisms such as niche differences or through competitors having similar relative fitness (19). Phage co-existence has previously been attributed to variation in host cell susceptibility to infection (20). In our system, ϕ14-1 and ϕLUZ19 use different bacterial surface receptors to infect cells (46,47). We propose that heterogeneity in the expression of phage receptors in the bacterial population (48) may have created a niche that enabled ϕLUZ19 persistence, but which was inaccessible to the rescued ϕ14-1 population. Alternatively, phage fitness differences may have been resolved through ϕLUZ19 co-evolution. For example, ϕLUZ19 acquired tail fiber and tail protein mutations which likely contribute to host attachment rates and within-host competitiveness (49). While the exact cause of co-existence remains unclear, the results highlight that stabilising mechanisms could buffer against total competitive exclusions that arise through evolutionary rescue.

Global biodiversity is decreasing due to environmental stress caused by land use change, pollution, and climate change (15,50,51). This study highlights a process by which evolutionary rescue, a force typically associated with preserving biodiversity, can make the community less resilient over ecological and evolutionary time. That parasites, and specifically viruses, can undergo evolutionary rescue has implications for our understanding of how parasites might evolve in the context of novel and hostile environments, such as following spillover events (52) or in hosts treated for infection (53). A loss of parasite diversity could increase the survival rates of some host species (54); there could be a reduced burden of infection, fewer co-infections, and weaker selection favouring virulence due to less inter-specific competition (55). Given parasites are beneficial for keeping pest or pathogenic hosts (as in this study) at bay (53,56), lower diversity would have negative consequences for animal, plant, and ecosystem health. Ultimately, consideration of the eco-evolutionary dynamics will help us better understand how communities will respond to increasingly frequent environmental stressors in a changing world.

## Methods and Materials

### Strains, storage, and culture conditions

*Pseudomonas aeruginosa* PAO1 (hereafter referred to as PAO1) was used with two lytic bacteriophages: ϕLUZ19 (46,57), and ϕ14-1 (58). These phages have been used to test several ecological and evolutionary hypotheses at 37°C (32,59–62). The phages were also selected based on their distinct responses to temperature: ϕLUZ19 has high growth rates at 37°C and 42°C, while ϕ14-1 has restricted growth at 42°C (32). Bacterial stocks and phage lysates were prepared as in ref (32).

### Experimental evolution

A schematic of the experimental evolution is shown in Figure 1. Each of 15 evolutionary passages were made across three phage treatments (ϕLUZ19 and ϕ14-1 monocultures and co-culture) and two temperatures (37°C and 42°C). Each treatment consisted of six, independent replicate populations started from a single ancestral lysate.

Phage were propagated without shaking with a non-evolving ancestral PAO1 bacterial host. For the initial passage, ancestral phage lysates were diluted to 10^8^ PFU/ml and 300μl were added to 2.7ml 10^8^ CFU/ml bacterial culture in loose-lid 14ml falcon tubes. Phage co-culture lines were prepared by combining 150μl each of ϕLUZ19 and ϕ14-1 10^8^ PFU/ml stocks prior to mixing with bacteria. Phages were added at a 1:1 ratio to mimic a phage community that is already stable and at the maximum level of biodiversity possible with a two species system. The initial passage phage densities were ∼10^7^ PFU/ml resulting in a phage/bacteria ratio (multiplicity of infection, MOI) = ∼0.1. Bacterial culture densities were standardised using optical density (OD595) based on a CFU:OD standard curve (Fig S2). Following addition of bacterial cultures, tubes were incubated statically at 37°C or 42°C in circulating water baths for 8h.

After each passage, phage lysates were centrifuged at 3,095xg for 5 mins to pellet remaining bacterial cells. Phage lysates were then sterile-filtered using 0.2μm syringe filters into 2ml cryotubes and stored at 4°C. At the beginning of each passage, 2.7ml of fresh ancestral PAO1 was seeded with 300μl of the preceding passage’s filtered phage lysate.

### Phage quantification

Phage titres were determined via the double-layer overlay method (63) following the same protocol as in ref (32). Briefly, bacterial lawns were prepared by mixing 10mL of melted top agar with 300µL of a *P. aeruginosa* PAO1 overnight culture. Phage stocks were serially diluted, and 10µL was spotted onto bacterial lawns. After incubating plates for 6–8 h at 37°C, spots with the highest number of discernible plaques were counted. Top-agar bacterial lawns were seeded with ϕLUZ19-resistant and ϕ14-1-resistant PAO1 strains to quantify ϕ14-1 and ϕLUZ19 lines, respectively. Phage resistant PAO1 strains were generated by spotting high titre phage stocks (∼ 10^10^ PFU/ml) onto wild-type PAO1 top-agar lawns. Plates were then incubated for ∼48h or until colonies started to grow on top of phage clearance zones. Five colonies were picked for each phage and were re-streaked twice before being used to seed 10ml Luria-Bertani (LB) (Lennox) and grown statically at 37°C in 50ml falcon tubes. The absence of phage in resistant cultures was determined through sterile-filtration of the bacterial supernatant and spotting onto ancestral PAO1 bacterial lawns.

Phage resistance was confirmed through the absence of plaques when spotting high titre phage stocks onto lawns seeded with each resistant line. One resistant line was selected for each phage by spotting phage stocks of known concentration (based on wild-type PAO1 estimates) and selecting the line that had the closest plaque count, turbidity, and size. Phage-resistant PAO1 mutants underwent whole-genome long-read sequencing and variant calling (see DNA extraction and sequencing, and sequence analysis for methods). While both PAO1 mutants had multiple mutations compared to wild-type (Table S3), we identified mutations that were previously linked to phage resistance. ϕLUZ19-resistant PAO1 had a mutation in a GspL type-II secretion system protein (BlastP: 99.5% similarity and 100% query cover); Gsp gene mutations have previously shown to provide resistance to type-IV pilus dependent phages (64). ϕ14-1-resistant PAO1 had a mutation in a glycosyltransferase (BlastP: 100% similarity and query cover); glycosyltransferase mutations have been shown to provide resistance to LPS-dependent phages (65). To maintain comparability, both monoculture and co-culture lines were quantified using resistant PAO1 strains.

Changes in phage counts during the evolution experiment could reflect evolutionary changes in efficiency of plaque formation rather than changes in phage densities. The efficiencies of plaque formation of the passage 15 evolved and ancestral phage lines were found to be similar when tested on the ancestral and resistant PAO1 strains separately (Fig. S6). This outcome confirmed that phage counts reflected changing phage densities.

### Phage separation and concentration

Evolved phage populations needed to be concentrated and purified to separate co-cultures and create high titre stocks for phage population sequencing (Fig. 1). High titre, pure phage stocks were made using a selective confluent lysis double-layer overlay method. Briefly, bacteria-phage lawns were prepared by mixing 180μl of either ϕLUZ19-resistant or ϕ14-1-resistant PAO1 overnight cultures (approximately 3 x 10^8^ CFU/ml) with 30μl of phage lysate diluted to 10^8^ PFU/ml. Initial bacterial culture densities were ∼10^8^ CFU/ml and phage densities were ∼10^6^ PFU/ml phage, MOI = ∼0.01. Bacteria-phage mixtures were left at room temperature for ∼10 mins to allow phage adsorption after which 5ml of molten top agar (∼40°C) was added and agar was poured onto pre-filled LB-agar plates. Plates were incubated for ∼20h at temperatures appropriate for each evolved population: 37°C for 37°C evolved phage populations, 42°C for 42°C evolved populations. After incubation, top-agar was scraped off plates into 15ml falcon tubes containing 5ml phage buffer (NaCl (100 mM), MgSO_4_ (10 mM), CaCl_2_ (5 mM), Tris-HCl (pH 8) (50 mM), Gelatin (0.01%)). Tubes were incubated at 4°C on a rotating carousel shaker at 10rpm for 24h to extract phage from top agar. Phages were separated from bacteria and agar by centrifuging tubes at 6,000xg for 10 mins followed by sterile-filtering. The purification/concentration process was repeated three times to remove non-focal phages and purity was assessed based on the absence of competitor plaques following high-titre spotting. To ensure comparability, monoculture lines underwent the same purification and concentration process as co-culture lines.

### Phage phenotypic assays

#### Growth rates

The phenotypes of purified evolved phage lines relative to the ancestor was assessed by measuring phage and bacterial growth across an 8h window under static incubation at 37°C and 42°C. Phage stocks were diluted to 10^5^ PFU/ml and 300uL was used to inoculate 2.7ml of 10^8^ CFU/ml wild-type PAO1, with a resulting MOI = ∼0.0001. This low MOI was chosen to extend phage growth curves to capture differences in phage growth rates as, at higher phage densities, both phages tend reach carrying capacity within 2-3h (32). For ϕLUZ19, samples were taken for phage quantification at 2h, 4h, and 8h. For ϕ14-1, samples were taken at 4h and 8h as preliminary assays indicated that phage growth was minimal at the 2h timepoint. Phage quantification was performed by adding 200μl samples to 96-well filter plates (Agilent) followed by centrifugation at 2,230xg for 5 mins before spotting onto ϕ14-1 or ϕLUZ19 resistant PAO1 double-layer overlay plates. Each fitness assay included a single replicate of each evolved phage line and three replicates of the phage ancestor. Growth rate assays were repeated three times across a two-week period to produce three technical replicates.

#### Competitive ability

We assessed the competitiveness of evolved 14-1 37°C and 42°C co-culture populations across time. Competitive ability was determined by growing phages under the same conditions as the fitness assay (37°C and 42°C) either alone or in the presence of an ancestral ϕLUZ19 competitor. For the monoculture treatment, 300μl of phage lysate was added to 2.7ml 10^8^ CFU/ml wild-type PAO1 stock. For the co-culture treatment, 150μl of evolved phage stock and 150μl of ancestral phage competitor was added. A 1:1 phage ratio was used to replicate the experimental evolution selective environment. The competition assay was conducted with two phage starting densities, 10^5^ PFU/ml (MOI = ∼0.0001) and 5 x 10^8^ PFU/ml (MOI = ∼5).

Phages were grown for 8h after which samples were taken for phage quantification as previously described. Evolved ϕ14-1 competitiveness was determined by calculating ancestral ϕLUZ19 competitor growth in co-culture with evolved and ancestral ϕ14-1 populations against phage growth in monoculture (66). Competition assays were repeated three times across a four-week period to produce three technical replicates.

### Phage population genomics

#### DNA extraction and sequencing

Phage DNA was extracted using a customised protocol. We used 500μl aliquots of post-purification evolved and ancestral phage lysates (∼10^10^ PFU/ml). Firstly, we added DNase (5μl of 1000U/ml, 5U) to remove bacterial DNA and RNase (2μl of 100mg/ml, 0.2mg) to remove RNA. Lysates were then incubated at 37°C in a heat block for 1h and inverted every 15 mins. After incubation, 67.5μl lysis (AL) buffer and 4μl proteinase K was added to each tube before incubating at 56°C for 15 mins. After 15 mins, tubes were then incubated at 95°C for 10 minutes to denature the proteinase K. After denaturing, the tubes were placed on ice and 150μl of precipitation (N4) buffer was added. Tubes were immediately centrifuged at 13,000xg for 10 mins to pellet cell debris and the supernatant was transferred to a clean 2ml Eppendorf. Cold 100% isopropanol (1.5x tube volume, ∼1.1ml) was then added and the tubes were placed in an orbital rotator set to 10rpm for 5 mins to precipitate DNA. The tubes were centrifuged at 13,000xg for 20 mins to pellet DNA after which the supernatant was discarded. The DNA pellet was then washed with 1ml 70% ethanol and mixed for a further 5 mins on the orbital rotator before being centrifuged again at 13,000xg for 20 mins. This wash step was repeated twice and after the second centrifugation step the supernatant was discarded and the pellet dried in a heat block set to 37°C to evaporate any remaining ethanol. Finally, 30μl of nuclease-free water was added to re-suspend the pellet.

DNA purity and contamination were measured using NanoDrop 2000c (Thermo Scientific). The presence of phage DNA was confirmed using gel electrophoresis using a 100kb ladder and a phage lambda DNA control with bands observed at the expected phage genome size. DNA was quantified using Qubit 4 (Thermofisher). Samples were diluted to DNA concentration of 50ng/μl and sent for Illumina short-read sequencing with AZENTA/GENEWIZ using their Microbe-EZ pipeline. One extraction from each evolved population was sent for sequencing in addition to three extractions of each phage ancestor. For ancestral and phage-resistant PAO1 bacterial genome sequencing, bacterial samples were sent to MicrobesNG for DNA extraction and long-read sequencing.

#### Sequence analysis

Phage sequence reads were pre-processed through read trimming using Trim Galore (v.0.5.0) (https://github.com/FelixKrueger/TrimGalore) with a fastqc step and 33 phred-score read cut off. Due to high and uneven read depth, reads were downsampled using bbnorm from the bbmap package (v.39.18) (https://sourceforge.net/projects/bbmap/) to a target read depth of 1500x and a minimum depth of 1000x. Ancestral phage genomes were assembled using shovill (v1.1.0) (https://github.com/tseemann/shovill) with default parameters and downsampled phage reads were mapped to the assemblies using Bowtie2 (v.2.3.4.2) (67) with default parameters. Read depth was checked for evenness using SAMtools (v.0.1.2) (68) view, sort, and depth functions. Ancestral phage assemblies were annotated using prokka (v.1.14.5) (69), guided by the NCBI GenBank file for each phage (ϕ14-1: NC_011703; ϕLUZ19: NC_010326). Genetic variants in phage lines were detected using breseq (v.0.36.1) (70) under default parameters, using the annotated ancestral genomes as a reference.

Wild-type PAO1 reads were processed using an in-house pipeline. We first quality-controlled the long-reads using Filtlong (v. 0.2.1) (https://github.com/rrwick/Filtlong) with parameters - -min_length 1000 --keep_percent 95. We then used Autocycler (v. 0.4.0) (71) to recover a consensus genome assembly, calling the assemblers Canu (v. 2.3) (72), Flye (v. 2.95), Miniasm (v. 0.3) (73), plassembler (v. 1.7.0) (74) and Raven (v. 1.8.3) (75). Next, we quality-controlled the short-reads using fastp (v. 0.24.2) (76), indexed with assembly with BWA (v. 0.7.19) (77), and polished with Polypolish (v. 0.6.0) (78). Lastly, we re-oriented the assembly with Dnaapler (v. 1.2.0) (79). The workflow was deployed using a Dockerised Nextflow pipeline (v. 1.0.2) available at https://doi.org/10.5281/zenodo.15706447. The wild-type PAO1 assembly was annotated using prokka (v.1.14.5) (69). ϕLUZ19 and ϕ14-1-resistant PAO1 mutations were detected by mapping long reads to the wild-type assembly with minimap2 (v.2.24) (80) and variant calling with medaka (v.2.1) (https://github.com/nanoporetech/medaka). SNPs were filtered so only those with quality scores >= 10 were kept.

### Statistics and data visualisation

All statistical analyses and data visualisation were conducted using packages in R (v.4.3.2) and RStudio (81,82). Data wrangling was performed using “Tidyverse” (v.2.0.0) R packages (83). Phage growth and evolution rates were compared between lines using linear mixed effect models with the “lme4” (v.1.1-36) R package (84) where the response variable was phage density (pfu/ml) or genetic distance from ancestor, the explanatory variable was an interaction term between evolution treatment and temperature, and batch was a random effect. Genetic divergence between evolved populations was calculated using Principle Coordinate Analysis (PCoA) ANOSIM with 10,000 permutations using the “ape” (v.5.8) R package (85). Congruence between Euclidean genetic and phenotypic distance neighbour-joining trees was calculated using Procrustes Approach to Cophylogenetic Analysis (PACo) (v.0.4.2) R package with 10,000 permutations (86). Data and code used in analyses can be found at https://github.com/SamuelGreenrod/Phage_thermal_adaptation.

## Acknowledgments

We thank R. Salguero-Gomez, T. Richards, T. Barraclough, and K. Foster for feedback on the experimental design and results. We also thank M. Whitlock, A. Hasan, R. Germain and D. Burstein for feedback on the manuscript, and M. Blazanin for feedback on analysing competition data. This work was supported by the Biotechnology and Biosciences Research Council (BB/T008784/1) to S.T.E.G. as well as the Natural Environment Research Council (NE/X000540/1) and NSERC Canada Excellence Research Chair to K.C.K. The funders had no role in study design, data collection and interpretation, or the decision to submit the work for publication. Phage sequence reads are accessible on NCBI (https://www.ncbi.nlm.nih.gov/) under BioProject ID: PRJNA1332698. Bacterial sequence reads are available on NCBI under BioProject ID: PRJNA1332799.

## Extended Data

**Figure S1.**
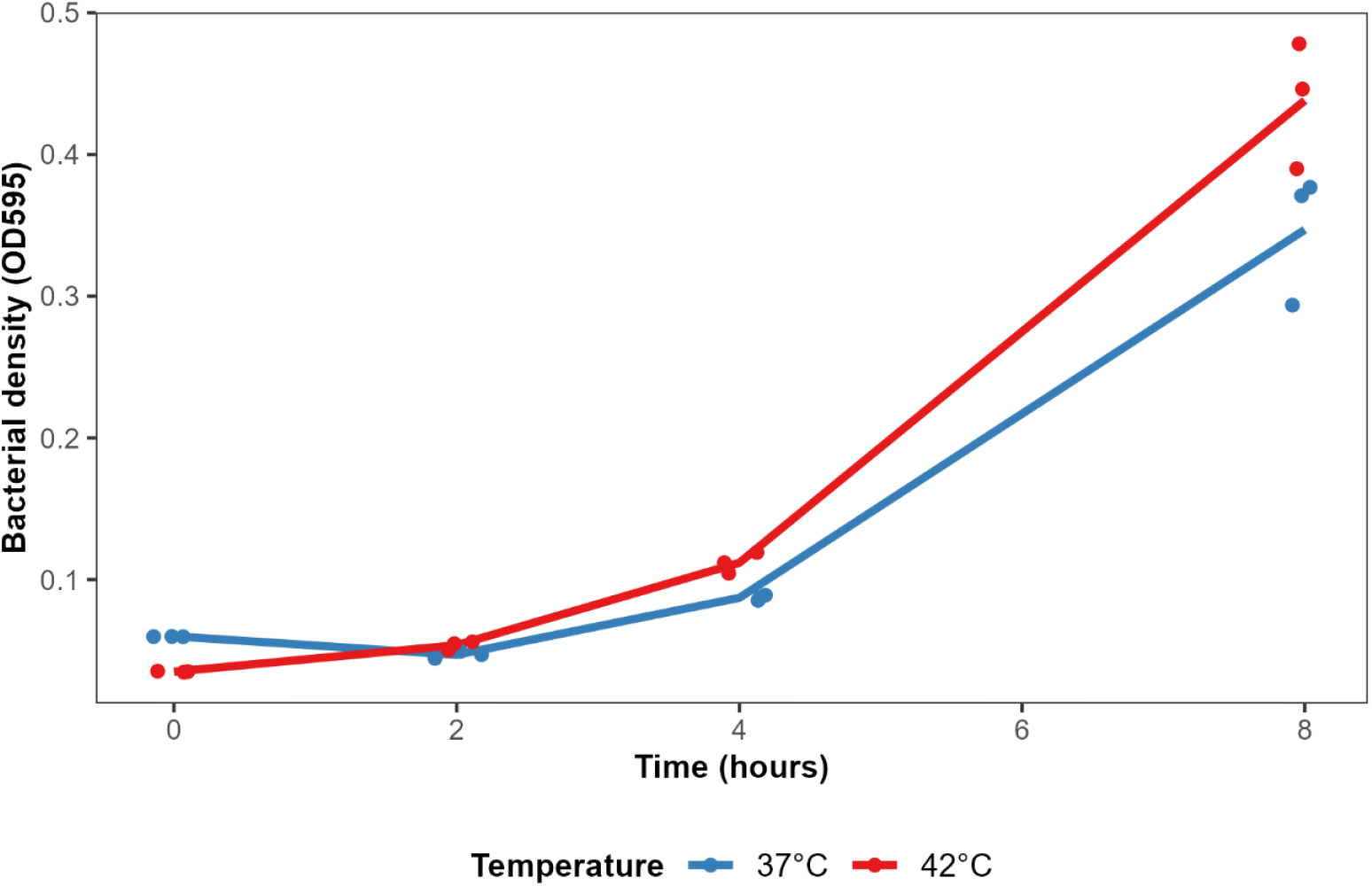
Bacterial growth in absence of phage is similar across temperature. Growth curves of no phage bacterial control. Dots reflect an average of three technical replicates. Bacterial growth was measured three separate times.

**Figure S2.**
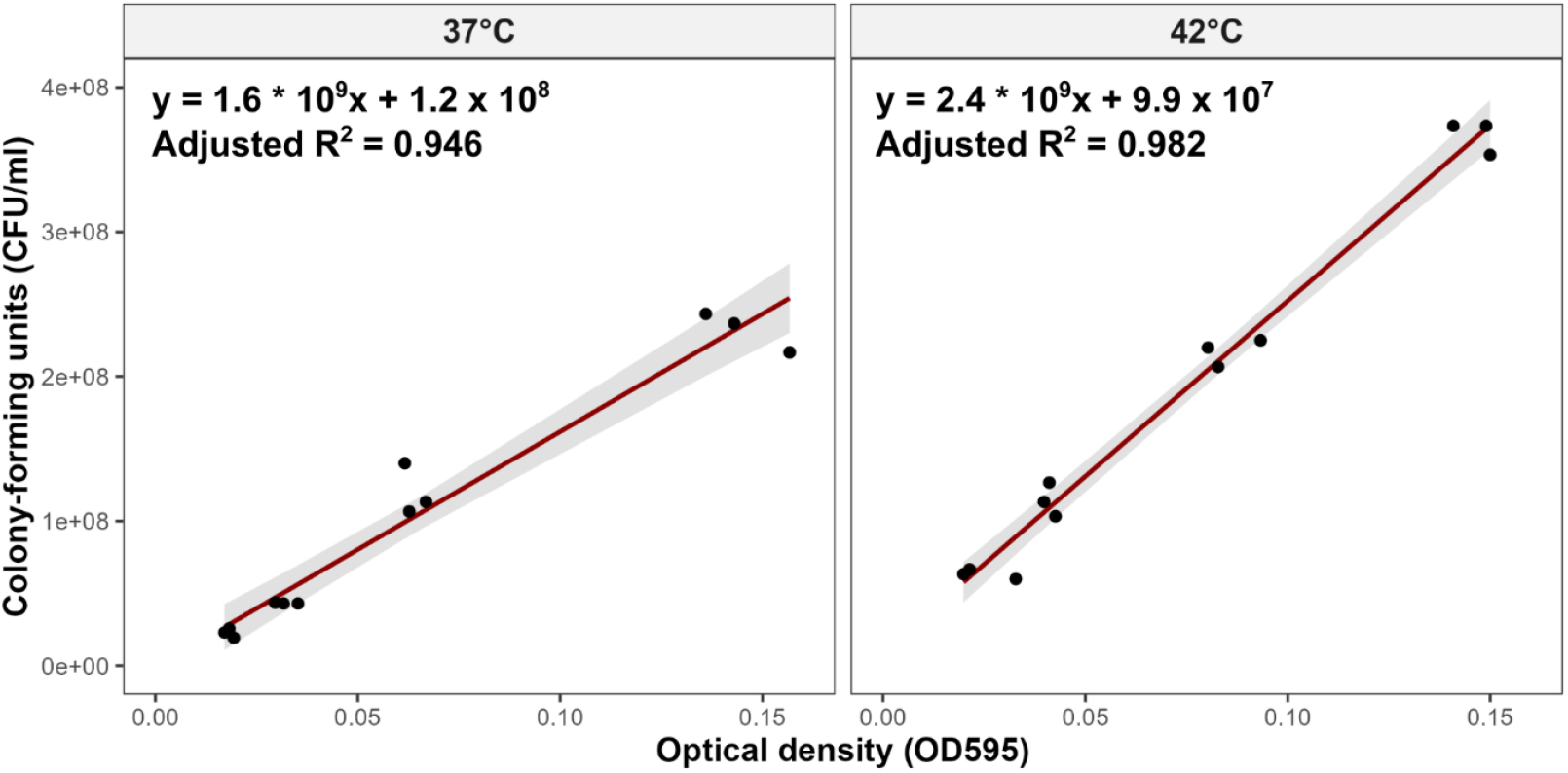
Optical density v colony-forming units standard curve. A regression of optical density and colony-forming units was used to determine the optical density of *P. aeruginosa* PAO1 overnights required to have ∼10^8^ CFU/ml.

**Figure S3.**
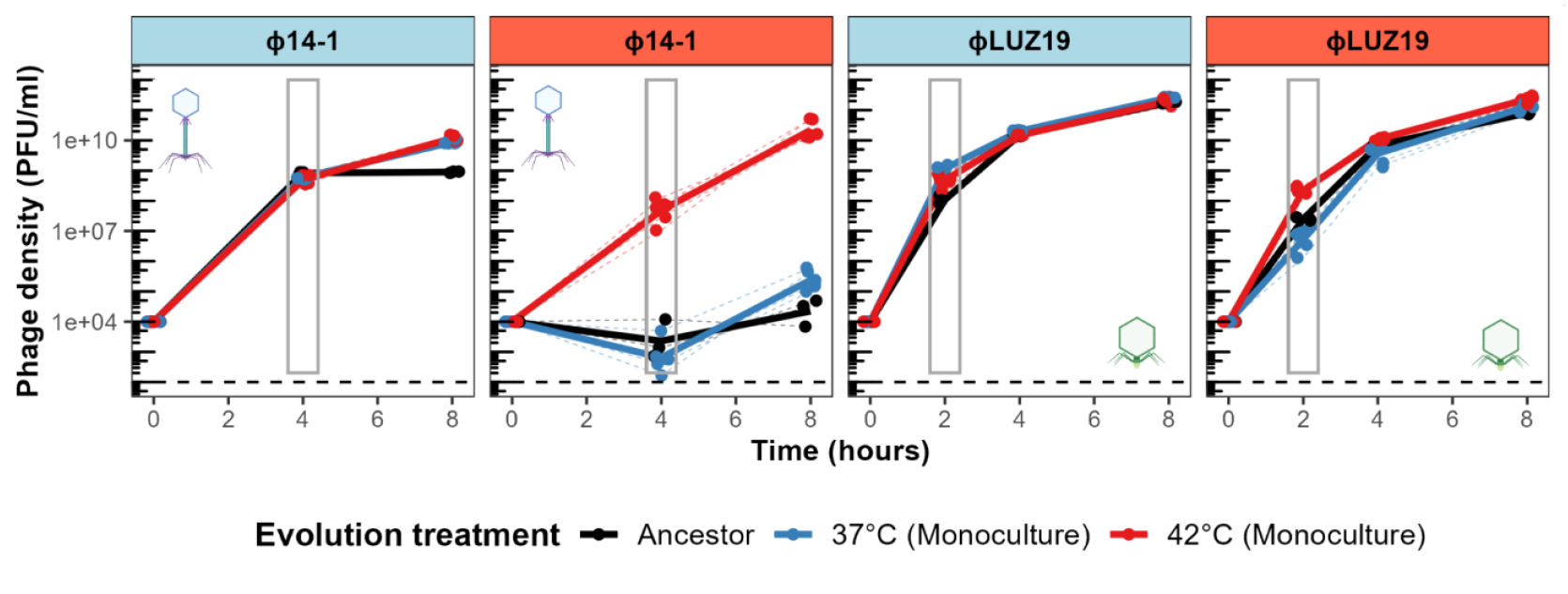
Evolved and ancestral phage growth curves at 37°C and 42°C. Grey box shows time points selected for fitness comparison. Black dashed line shows lower detection limit. Phage growth was assessed at 37°C (light blue strip) and 42°C (light red strip).

**Figure S4.**
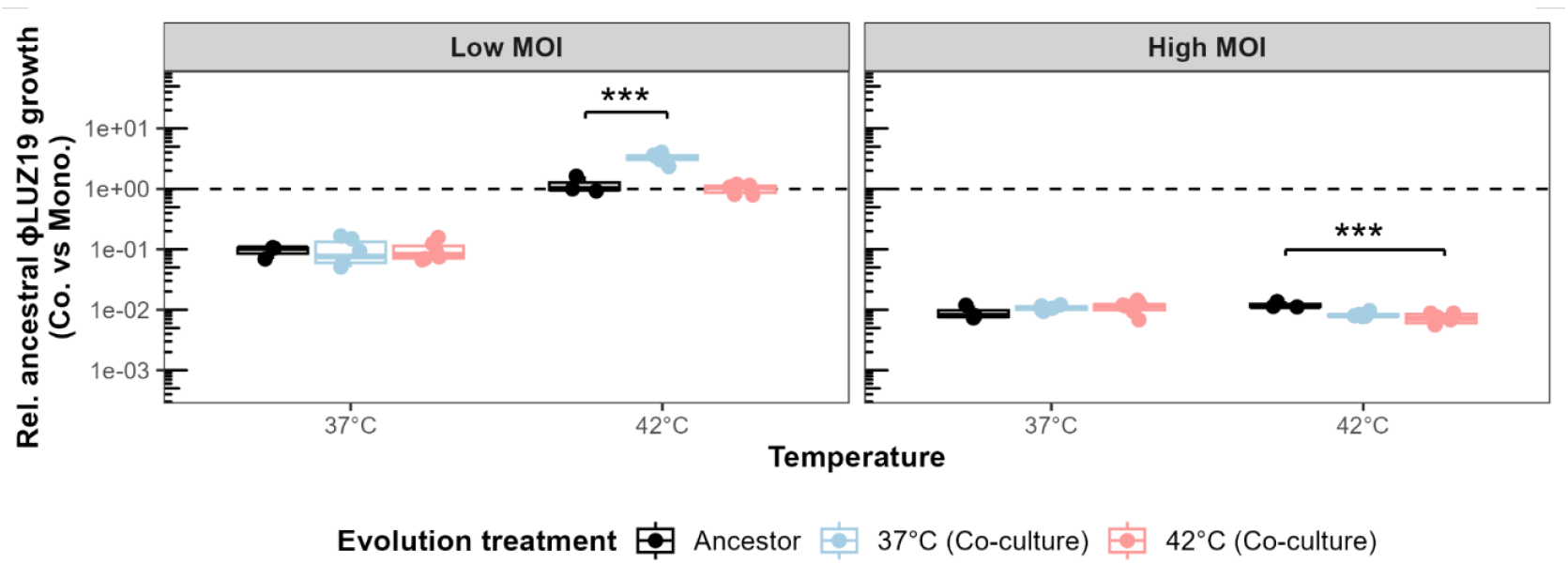
ϕ14-1 co-culture evolved populations have similar competitiveness to ancestor. Boxplots show the growth of ancestral ϕLUZ19 at 37°C and 42°C in co-culture with the ancestral and co-culture evolved ϕ14-1 populations relative to growth in monoculture. ϕ14-1 ancestral and evolved population competitiveness was determined at both low MOI (MOI = 0.0001) and high MOI (MOI = 5). Values below 1 (dashed black line) indicate ϕLUZ19 growth is inhibited by the presence of ϕ14-1 populations. Asterisks show significant differences in growth restriction between ϕ14-1 evolution treatments. ** = p < 0.01.

**Figure S5.**
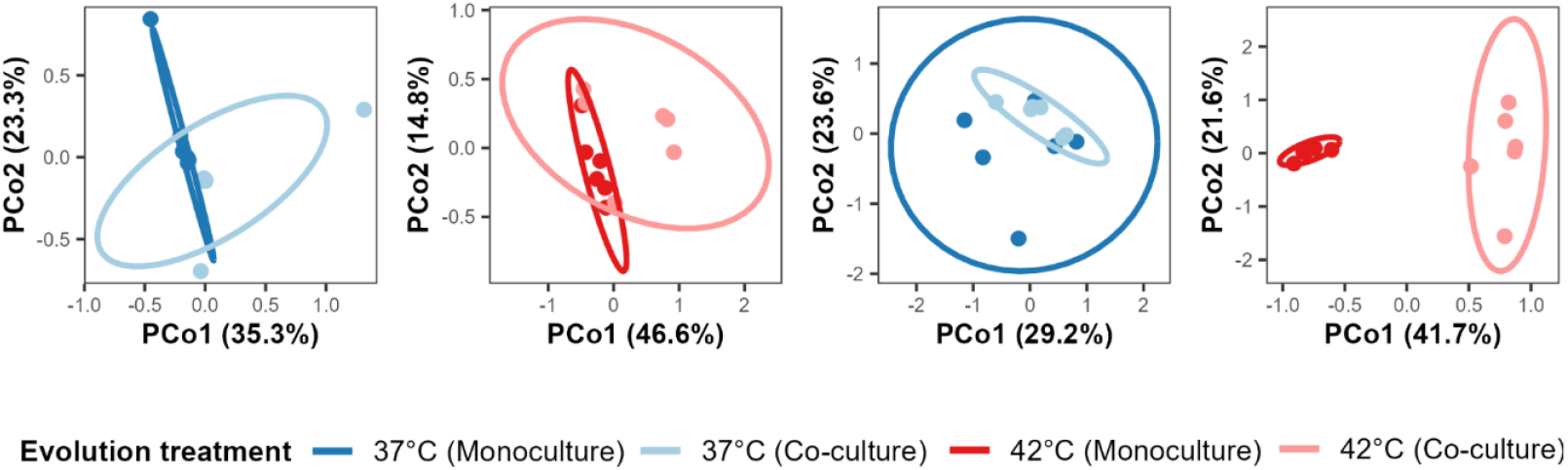
Phages show significant genetic divergence between monoculture and co-culture populations. PCoA plots show Euclidean genetic distance clustering between monoculture (37°C in deep blue, 42°C in deep red) and co-culture (37°C in light blue, 42°C in light red) evolved lines.

**Figure S6.**
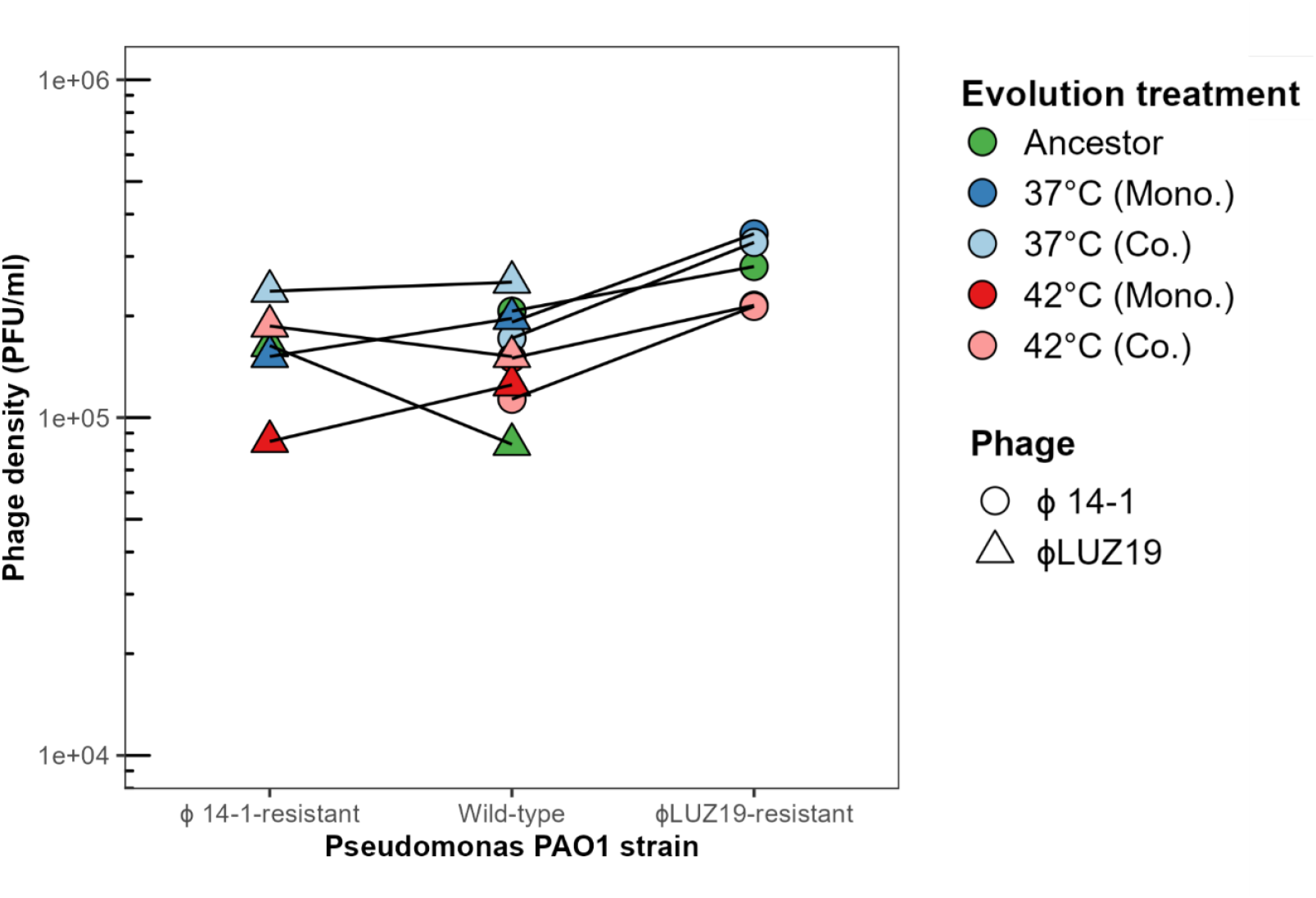
Efficiency of plaque formation is the same for evolved and ancestral phage lines. Plot shows the density of evolved and ancestral phage lysates as measured with plaque assays on the wild-type and resistant PAO1 strains. Lines between points show phage counts on the same stocks on separate hosts.

